# The development of the relationship between auditory and visual neural sensitivity and autonomic arousal from 6m to 12m

**DOI:** 10.1101/2022.12.15.520575

**Authors:** K. Daubney, Z. Suata, I. Marriott Haresign, M.S.C. Thomas, E. Kushnerenko, S.V. Wass

## Abstract

The differential sensitivity hypothesis argues that environmental sensitivity has the bivalent effect of predisposing individuals to both the risk-inducing and development-enhancing influences of early social environments,. However, the hypothesis requires that this variation in environmental sensitivity be general across domains. In this study, we focused on neural sensitivity and autonomic arousal to test domain generality. Neural sensitivity can be assessed by correlating measures of perceptual sensitivity, as indexed by event-related potentials (ERP) in electrophysiology. The sensitivity of autonomic arousal can be tested via heart rate changes. Domain generality was tested by comparing associations in perceptual sensitivity across auditory and visual domains, and associations between sensitivity in sensory domains and heart rate We contrasted ERP components in auditory (P3) and visual (P1, N290 and P4) detection-of-difference tasks for N=68 infants longitudinally at 6 and 12 months of age. Domain generality should produce correlated individual differences in sensitivity across the two modalities, with higher levels of autonomic arousal associating with increased perceptual sensitivity. Having controlled for multiple comparisons, at 6 months of age, the difference in amplitude of the P3 component evoked in response to standard and deviant tones correlated with the difference in amplitude of the P1 N290 and P4 face-sensitive components evoked in response to fearful and neutral faces. However, this correlation was not found at 12 months of age. Similarly, autonomic arousal negatively correlated with neural sensitivity at 6 months but not at 12 months. The results suggest neural perceptual sensitivity is domain-general across auditory and visual domains, and is related to autonomic arousal at 6 months but not at 12 months of age. We interpret these findings within a neuroconstructivist framework and with respect to the concept of interactive specialisation. By 12 months of age, more experience of visual processing may have led to top-down endogenous attention mechanisms that process visual information in a way that no longer associates with auditory perceptual sensitivity.

---

Individuals vary systematically in their sensitivity or “permeability” to experiential and contextual influences on development and health (Boyce, 2015). Environmental Sensitivity (ES) theorists posit that there is a common factor of sensitivity along which individuals differ in their ability to register and process environmental stimuli (Pluess, 2015). Those who are especially sensitive are unusually susceptible not only to the risk-inducing but also to the development-enhancing influences of early social environments (Belsky et al., 2007; Boyce & Ellis, 2005; Ellis et al., 2011).

But what exactly does it mean, mechanistically, for one individual to be more sensitive than another to both risk-inducing and development-enhancing influences? Within the field of ES, a wide range of traits have been used to index sensitivity that can be categorised into genetic (polygenic risk scores (Nelemans et al., 2021)), physiological (e.g., cortisol reactivity (Obradović et al., 2010), autonomic nervous system activity (Weyn et al., 2022)) and behavioural/psychological sensitivity factors (e.g., negative emotionality (Kim & Kochanska, 2012) (For a review see Belsky & Pluess, 2013). Much research is based on reporting cross-over interactions where the effect of a positive (maternal empathy (Pitzer et al., 2011)) or negative (maternal depression (Netsi et al., 2015; Sacchi et al., 2018)) contextual measure on some behavioural outcome (infant sleep (Netsi et al., 2015), infant motor activity (Sacchi et al., 2018), later externalising problems (Pitzer et al., 2011)) is moderated by the value of a sensitivity measure. This provides support for the bivalency of sensitivity propounded by ES theorists. However, studies that look specifically at only one index of sensitivity are not able to address whether an individual’s environmental sensitivity varies across different levels of measurement or whether differences in sensitivity in all domains covary (Pluess, 2015; Stamps, 2016). Theoretically all measures of environmental sensitivity should correlate for it to be a domain-general trait. In addition, to our knowledge, no previous research has looked at how different domains work in concert in infants to further understand how different phenotypical markers of sensitivity develop. This study considers two branches, one considering correlations between perceptual modalities, the other from perceptual modalities to autonomic arousal, all being possible proximal measures of environmental sensitivity.

This paper set out to examine whether there were associations between envirionmental sensitivity (ES) as previously operationalized in studies looking at visual perception and auditory perception, and also to examine how these associations develop over the first 12 months of life. Several researchers have suggested that domain-general neural mechanisms may become more domain-specific over the course of development - including ideas relating selectionism to synaptic pruning (Kerszberg et al., 1992), hypotheses on the increasing localization of language areas (Neville et al., 1992), the “perceptual narrowing” hypothesis (Scott et al., 2007), and proposals about the increasing restriction of functional cortical areas with development (Durston et al., 2006). Interactive Specialization is also situated within a broader context of work on “neuroconstructivism” (Elman et al., 1996; Mareschal et al., 2007; Karmiloff-Smith, 2009). This corroborates the idea that early in development, an initial broad multisensory perceptual tuning, and the relative lack of cross-modal interactions, means that young infants do not integrate low-level features of sensory information into a single percept, as they do later in development (Boothe, 2010), and instead process input from auditory and visual modalities separately and in parallel (Lewkowicz & Ghazanfar, 2009)

Event-related potentials (ERPs) evoked in response to external stimuli have been used to chart the development of automatic-attentional processes (Kushnerenko et al., 2002). Neural markers of automatic auditory perception can be induced using oddball paradigms where frequently presented ‘standard’ tones are interspersed with less frequent ‘deviant’ tones. Recording deviance-elicited brain responses using EEG is a feasible way to assess automatic auditory discrimination and regularity detection abilities in even very young infants (Kushnerenko et al., 2013). The mismatch response (MMR) is a neurophysiological indicator of automatic, pre-attentive change detection between consecutive sounds and heightened sensitivity to deviant stimulus (Näatänen & Alho, 1995; Wetzel & Schröger, 2014). In infants younger than 12 months of age the MMR is often found as a positive deflection between 150 and 300ms post change onset (Morr et al., 2002; Garcia-Sierra et al., 2011; Kushnerenko et al., 2013). One way to interpret individual differences in ERP amplitude is in terms of differences in distractibility. This is because a (positive) deflection at this latency means the MMR can merge/overlap with the P3a, which is generally understood to be the central electrophysiological marker of involuntary attentional orienting to a novel or unexpected sound (Friedman et al., 2001; Squires et al., 1975). It indexes involuntary (bottom-up, saliency driven) attention mechanisms (Escera et al., 2000; Friedman et al., 2001). This automatic orienting and attentional capture could be interpreted as less automatic inhibition of response (Kushnerenko, 2002) and therefore greater automatic neural sensitivity to environmental effects (Wass et al., 2018).

Another common indicator of involuntary neural sensitivity is neural responsiveness during emotional-face processing. Research investigating the ontogeny of the processing of faces in infancy has shown that between 5–7 months of age, ERP components associated with infant perceptual sensitivity to faces (occipitotemporal P1, N290 and P4 components) begin to reliably differ between fearful and non-fearful facial expressions: In 7-month-old infants the P4 was larger in response to fearful than neutral or happy faces (Leppänen et al., 2007); 7-month old infants had a larger P4 for fearful than angry faces (Kobiella et al., 2007); the amplitudes of the P1, N290 and P4 differed for fearful and happy faces in 7-month old infants (Nelson et al., 1996); 7-month old infants rated higher in perceptual sensitivity had larger N290 responses to fearful than to happy faces (Jessen & Grossmann, 2015); by 7-months of age, the N290 was larger in response to fearful than to happy, neutral or phase-scrambled faces (Yrttiaho et al., 2014). In addition, neural responses of 3-month-olds to fearful and happy faces differed over occiptotemporal regions implicated in face perception but not frontocentral regions implicated in attention. Early perceptual sensitivity may presage the attentional bias for fearful faces (Safar & Moulson, 2020).

In terms of autonomic correlates of sensitivity, in infants, higher heart rate (HR) has been found to associate with hypervigilance (Mammen et al., 2017). Associations between autonomic activity and sensory perception are largely limited to behavioural markers such that increased autonomic arousal associates with decreased voluntary attention control and increased responsivity to salient targets (Alexander et al., 2007; Arnsten, 2009; Liston et al., 2009). Only recently have researchers looked at how neural sensitivity, measured in terms of involuntary auditory attention using an auditory oddball task, varies with levels of autonomic arousal (Wass et al., 2019). They found that 5-7-year-old children with higher autonomic arousal showed larger P150/P3a amplitudes in response to small acoustic contrasts (500Hz-750Hz). This supported the notion that higher autonomic arousal associated with less inhibition of response to exogenous stimuli, which meant that even small acoustic contrasts could potentially elicit a P3a-like response.

The current study collected ERP data from infants presented with an auditory-oddball paradigm and a visual emotional faces paradigm. We examined whether neural sensitivity to auditory and visual stimuli were correlated over temporal and occipital regions respectively implying domain general sensitivity, or uncorrelated, implying that neural sensitivity is domain specific. We also examined how this domain specificity or generality changed between 6m and 12m. We also examined the relationship between neural sensitivity and autonomic arousal. Based on previous findings, we predicted that in order to support the theory that Environmental Sensitivity is domain general, increased autonomic arousal should associate with heightened neural sensitivity to differences in auditory stimuli as well as differences in visual stimuli.

## Method

### Participants

Infant-parent dyads attended the BabyLab at the University of East London on two occasions – first when the infants were 6m old and a second visit when the infant was 12m old. The participating parent-infant dyads were recruited from local children’s centres, baby sensory classes and new-parent support groups. Parents gave informed consent prior to the commencement of data collection.

#### Participant exclusions

At phase-one, 82 typically developing infants, (male 42 female 40), with a mean (*SD*) age of 27.5 (*2.4*) weeks on the day of testing, attended.

##### EEG – 6m

Data from a number of participants at phase one were unavailable due either to insufficiently good quality recording from one of the measures (designated so after visual inspection of the raw data and referral to video and session notes on the affective state of the infant during the recording) and were dropped before being processed (N=6), or fewer than 70 percent of the maximum number of trials in each condition trials on which to base the analysis (Monroy et al., 2021) (N=8 and N=16 for the auditory oddball and emotional faces paradigms, respectively). In total EEG data were available for N=68 and N=60 participants for the auditory oddball and emotional faces paradigms, respectively.

##### ECG – 6m

Insufficiently good ECG data (designated so after visual inspection of the raw data when the analysis software had identified almost the entire recording as noisy based on the default noise detection level of medium) led to a loss of data from N=8 participants. ECG data were available for N=74 participants; both ECG and EEG data were available from N = 60 and N=55 for the auditory oddball/emotional faces tasks respectively. The average age (*SD*) of participants who contributed both usable ECG and EEG faces data was 27.08 (*2.23*) weeks on the day of testing.

At phase-two, 68 of the initial cohort of 82 babies returned (male 36 female 32) with a mean (*SD*) age of 53.03 (*3.04*) weeks on the day of testing. Insufficiently good quality EEG data led to the loss of data from N=12 participants (see above). Insufficiently good ECG data (see above) led to a loss of data from N=5 participants. After pre-processing, participants were excluded due to not reaching the inclusion threshold for minimum numbers of trials (fewer than 60 percent of the maximum number of trials in each condition) for the EEG auditory oddball data N=5 and for the faces data: N = 7. In total, EEG auditory-oddball data were available for N=49 participants and EEG faces data were available for N=47 participants; ECG data were available for N = 63 participants; both ECG and EEG data were available from N = 46. The average (*SD*) of participants who contributed both usable ECG and EEG data on the second visit was 53.8 (*2.99*) weeks on the day of testing.

### Equipment

EEG was recorded using a high-density 128-channel HydroCel Geodesic Sensor Net (HGSN) produced by EGI (EGI, Eugene, OR). The EEG signal was referenced to the vertex, recorded at a 500 Hz sampling rate with band-pass filters set from 0.1–100 Hz using a Kaiser Finite Impulse Response filter. Prior to recording, the impedance of each electrode was manually checked to ensure that they were below 100 kΩm. ECG was recorded using a BioPac (Santa Barbara, CA) system recording at 1000Hz. ECG was recorded using three disposable Ag–Cl electrodes, placed in a modified lead II position. Stimuli were presented using Matlab.

### Procedure

Infants were seated on parents’ laps and presented with two paradigms. These were presented in four blocks, each lasting about 60 seconds. In order to maintain engagement, the two tasks were presented interspersed with excerpts showing nursery rhymes sung the children’s TV entertainer Mr Tumble. If participants were engaged with stimuli and calm, testers would proceed straight to the next block without pausing. The total paradigm, including preparation, recording, breaks and EEG cap removal, lasted approximately 40 minutes per participant.

#### Auditory oddball paradigm

This consisted of 100 trials. Each block consisted of: 70 ‘standard’ 500Hz tones; 15 ‘deviant’ 750Hz tones; 15 ‘noise’ (broadband white-noise) segments. The intensity of the tone and white-noise sounds was 70 dB sound-pressure level (SPL). The harmonic tones of 500 and 750 Hz fundamental frequency were constructed from the three lowest partials, with the second and third partials having a lower intensity than the first one by 3 and 6 dB, respectively. The harmonic tones were used instead of sinusoids for two reasons. Firstly, because it has been shown previously that complex tones result in larger N250 amplitudes in children then sinusoids (Čeponiené et al., 2001). Secondly, because we aimed to use the same paradigm that was used in a number of longitudinal and cross-sectional studies in infants and children in order to increase our understanding of the previously observed effects (Kushnerenko et al., 2007).

The duration of all sounds was 100 ms, including 5-ms rise and 5-ms fall times. The interstimulus (offset-to-onset) interval was 700 ms. The order in which the trials were presented was pseudo-randomised in order to ensure that two deviant and noise trials were always separated by at least two standard trials.

#### Emotional face paradigm

This paradigm consisted of the neutral and fearful expressions of 12 young (under 30-years) women’s faces taken from the Nim Stim faces database (Tottenham et al., 2009). The faces were pseudo-randomised so that the same face did not appear more than twice consecutively. Both facial expressions –neutral and fearful - appeared 23 times (+/- 2) each per block. There were four blocks, making 92 trials of each facial expression in total. The reason that 12 different faces were chosen for this study was to provide a variety of ethnicities that would reflect the demographic spread of participating families. A fixation appeared on the screen for 1000ms followed by a face for 500ms (see Fig. 1. for example fixations and faces). This meant that the ISI between faces was 1000ms. Evidence suggests that the optimal ISI for infant engagement and sustained attention during stimulus presentation is 600– 1,000 ms, which increases the presentation complexity and provides sufficient time for information processing (Xie & Richards, 2016).

**Fig.1.**
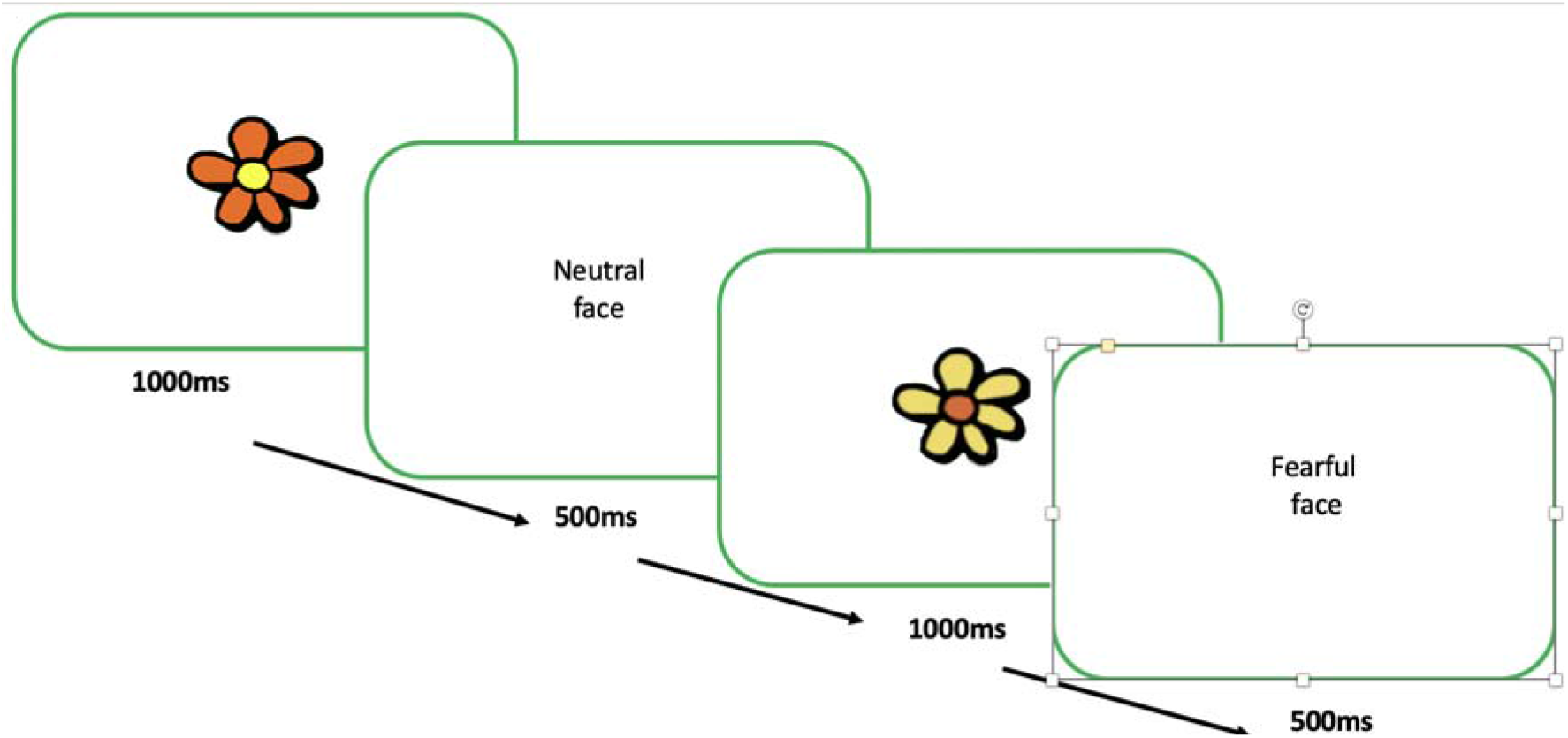
presentation sequence of stimuli for emotional face processing paradigm

### Data analysis

EEG data was processed using NetStation software (version 5.4.2). The vertex-referenced EEG was algebraically recomputed to an average reference. The signal was low-pass filtered off-line at 30 Hz using a Kaiser Finite Impulse Response filter and segmented into epochs starting 100 ms before and ending 800 ms after the stimulus onset. Artifact detection settings identified bad channels as those in which the amplitude exceeded 200μV using a moving average of 80ms. With infant nets there are no horizontal or lower eye channels. Blink detection is performed on a moving average of 80ms from the upper eye-channel minus its inverse. The threshold for exclusion was 140uV. Channels were marked bad for the entire recording if bad for greater than 30 percent of segments. Trials were marked bad if they contained more than 30 bad channels. Trials in which the number of channels marked bad, minus the number of channels marked bad for the entire recording exceeds 30, were excluded

#### EEG - Auditory oddball task

##### Exclusions

At 6m, the mean (range) [SD] number of trials included was 224 (210–270) [19] for standard; 57 (42–56) [4] for deviant; 48 (48–60) [4] for noise; At 12m the mean (range) (SD) number of trials included was 254 (179–280) [19] for standard; 55 (40–60) [4] for deviant; 55 (41–60) [4] for noise. This number of accepted trials has proven to be sufficient for this type of paradigm (Kushnerenko et al., 2013a; Guiraud et al., 2011; DehaeneLambertz and Dehaene, 1994; Friederici et al., 2007; Kushnerenko et al., 2013b, 2008).

##### Extracting average amplitude and latency

The valid ERPs obtained for each stimulus type were first averaged to create a per-participant mean waveform. The average of fronto-central channels was used (24, 20, 13, 19, 12, 11, 6, 5, 4, 124, 118, 112 (see fig. 2. c)) as the largest MMR/P3 was expected to occur over this area (Gumenyuk et al., 2005, 2004) and because it corresponded to those used to analyse data collected using the same paradigm previously (Kushnerenko et al., 2007, 2002b; Wass et al., 2019). Epochs were baseline-corrected to the average amplitude in the 100ms pre-stimulus period. The grand average (GA) waveform showed a clear difference between the amplitude of the P3 component in response to standard and deviant tones (see fig. 2. a and b). Therefore, the mean amplitude of the ERP to standard tones between 200 and 400 ms post stimulus onset was subtracted from the mean amplitude of the ERP to deviant tones in the same window to create a difference score between standard and deviant tones. Analysing the difference wave within this time-window was also in line with longitudinal and cross-sectional research using the same paradigm. (Kushnerenko et al., 2007, 2002b; Wass et al., 2019). Average amplitude was chosen as the most objective way to compare values between the standard and deviant conditions (Luck, 2014). This is because the latency of the peak is variable and sometimes it is not possible to identify the peak at all in young infants.

**Fig. 2.**
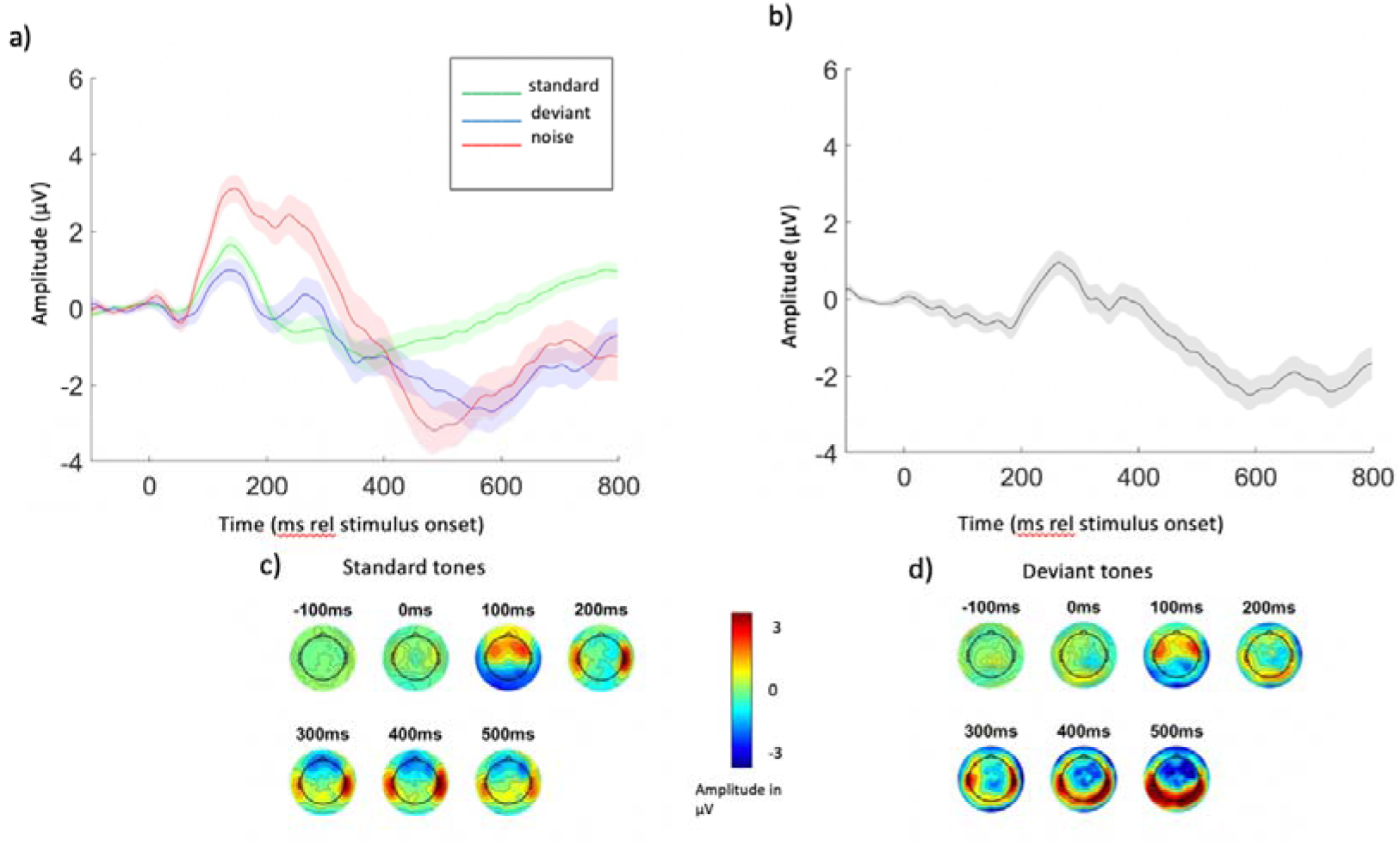
a) Grand average ERPs at 6m to auditory oddball task. Shaded areas represent the error bars, calculated as the Standard Error of the Mean b) Deviant-Standard difference wave (grand average). Shaded areas represent the error bars, calculated as the Standard Error of the Mean. c) Electrode locations used to calculate all ERPs. The locations used are marked in green. d) and e) Topoplots showing response to standard d) and deviant e) tones at 100ms intervals starting at 100ms pre-stimulus onset and ending 500ms post stimulus onset. Each topoplot shows an average of activity 50 ms around the given value (i.e. −100ms shows the average from −150ms to −50ms).

#### EEG – Emotional faces task

##### Exclusions

At 6m, the mean (range) [SD] number of trials included was 67 (42–91) [16] for neutral faces; 68 (48–91) [14] for fearful faces. At 12m the mean (range) [SD] number of trials included was 47 (39–59) [5] for neutral faces; 46 (40–58) [5] for fearful faces.

##### Extracting average amplitude and latency

The Grand Average (GA) waveform showed clear P1, N290 and P4 components in response to both face conditions (see Fig. 3. a and b). Therefore, the mean amplitude of the response to neutral faces in windows corresponding to the components P1 and P4 (between 50 and 150 ms and 350 and 450ms post stimulus onset) was subtracted from the mean amplitude of the response to fearful faces in the same windows. As the N290 is a negative-going component the mean amplitude of the response to fearful faces was subtracted from the mean amplitude response to neutral faces in the window 250-350ms post stimulus onset to create a difference score reflecting the absolute size of the difference in amplitude response evoked by the two conditions for this negative component. The average of occipital channels was used (64, 58, 51, 52, 59, 65, 69, 53, 60, 66, 70, 61 67, 71, 62, 72, 75, 76, 77, 78, 83, 84, 85, 86, 89, 90, 91, 92, 95, 96, 97, 98 (see Fig. 4. c)) as the largest infant facial perception components (P1, N290 and P4) were expected to occur over this area for a 128-electrode EEG cap (Haan et al., 2002; Halit et al., 2003; Vogel et al., 2012; Leppanen et al., 2007)

**Fig. 3.**
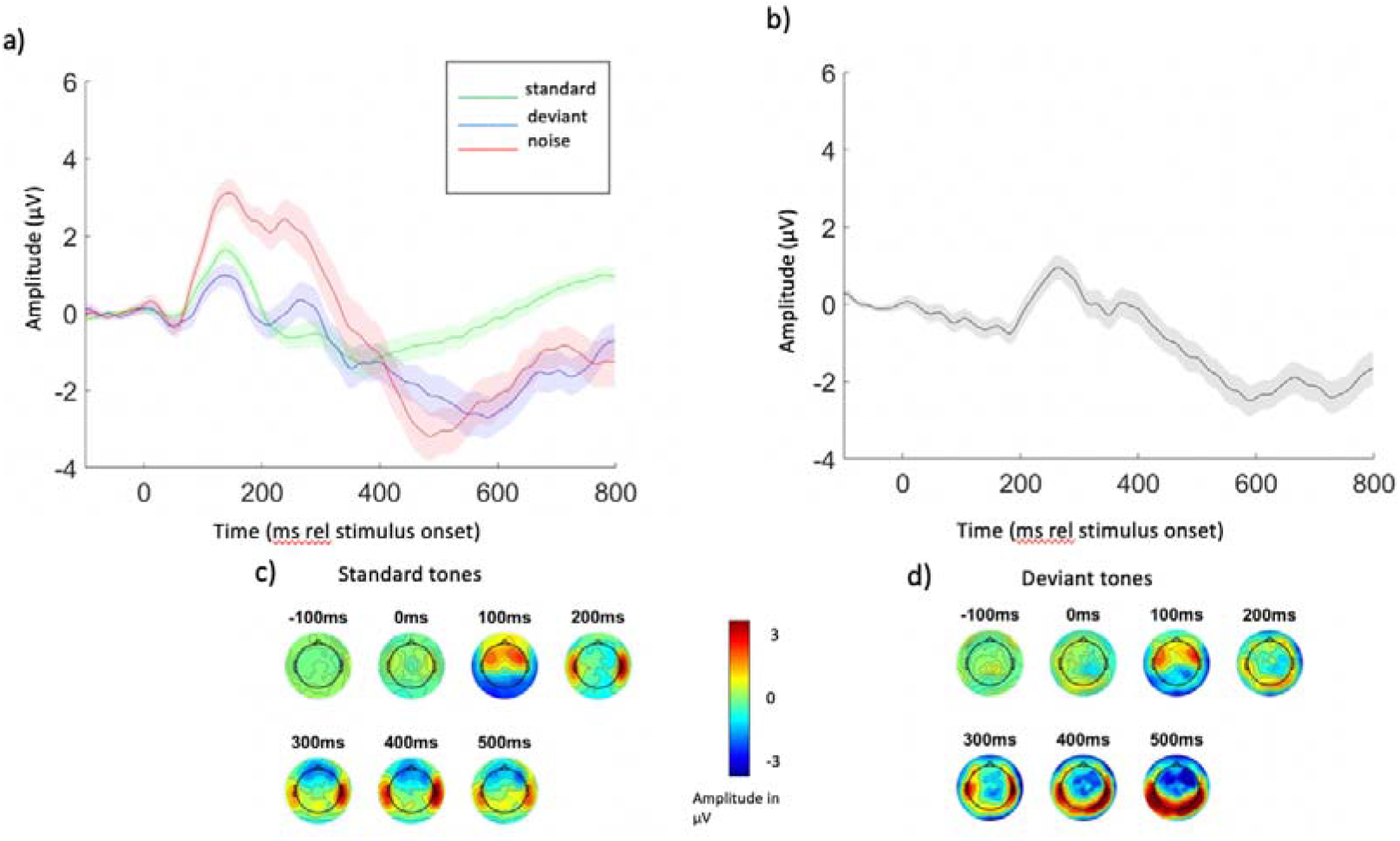
a) Grand average ERPs at 12m to auditory oddball task b) Deviant - Standard difference wave (grand average). Shaded areas represent the error bars, calculated as the Standard Error of the Mean c) and d) Topomaps showing response to standard c) and deviant d) tones at 100ms intervals starting at 100ms pre-stimulus onset and ending 500ms post stimulus onset. Each topoplot shows an average of activity 50 ms around the given value (i.e. −100ms shows the average from −150ms to −50ms).

**Fig. 4.**
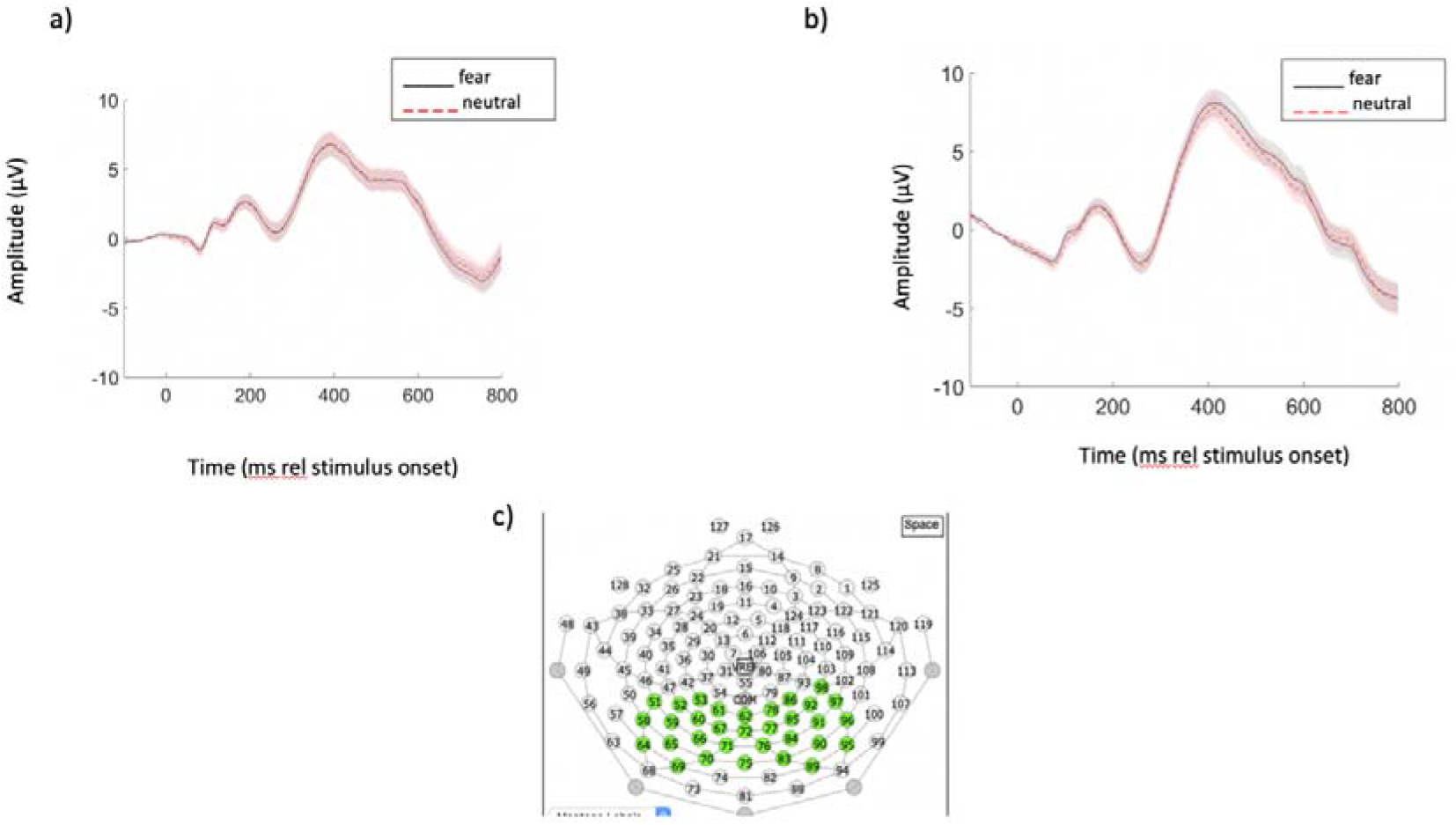
a) grand average ERPS to emotional faces at 6m. and b) 12m. Shaded areas represent the error bars, calculated as the Standard Error of the Mean c) Electrode locations used to calculate all ERPs. The locations used are marked in green.

#### ECG

Raw ECG data were analysed using Kubios software (Tarvainen et al., 2014). The R-wave time instants are automatically detected by applying the built-in QRS detection algorithm based on the Pan–Tompkins algorithm (Pan and Tompkins, 1985). The software automatically identified noise segments (using default setting of medium) based on the raw ECG data and from the interbeat interval data (RR or pulse-to-pulse intervals). Automatic artifact detection and rejection criteria were used to identify artifactual beats from the time series data consisting of differences between successive RR intervals and corrected in Kubios. The method has been validated (Lipponen and Tarvainen, 2019). Information on the algorithms used to process the raw ECG data in Kubios is included in supplementary materials. Heart rate was averaged across the duration of the recording while infants were presented with stimuli in order to replicate analyses using the same paradigm with 5-7-yr-old children (Wass et al., 2019).

### Statistical Analysis

After correcting for multiple comparisons using the Benjamini-Hochberg correction to control the false discovery rate, Bayesian statistics were used throughout. Bayesian statistics allow accepting and rejecting the null hypothesis to be put on an equal footing by providing a direct measure of the strength of evidence not only for but also against the study hypothesis, unlike frequentist statistical approaches, which do not determine whether non-significant results support a null hypothesis over a theory, or whether the data are just insensitive (Andraszewicz et al., 2015). Analyses were carried out using JASP software (Love et al., 2019). Bayesian Factor (BF) 10 gives the likelihood of the data under the alternative hypothesis divided by the likelihood of the data under the null so that BF10 values greater than 1 signal more confidence in rejecting the null hypothesis and values less than 1 signal more evidence in favour of the null. The BF01 is simply 1/BF10, that is, the likelihood of the data under the null compared to the alternative. The BFs above 1 indicate correlations for which the evidence from the current study is more likely under the hypothesis that there is a relationship between those variables in the population than not. A BF greater than 3 indicates “moderate” evidence for the study hypothesis that the two variables are correlated in the population. A BF greater than 10 indicates “strong” evidence for the study hypothesis that the two variables are correlated in the population. Prior and posterior information about the correlation summarizes how our knowledge about the unknown population correlation, in which all possible values from −1 to 1 were considered equally likely (prior), has changed as a result of information gathered in our study to put more weight on positive or negative values (posterior). (Nuzzo, 2017).

## Results

In Analysis 1 we examine the relationship between auditory and visual neural sensitivity at 6m and 12m. In Analysis 2 we examine the relationship between autonomic states and neural sensitivity at 6m and 12m.

### Preliminary analyses – descriptive

#### Auditory task

Fig. 2. shows the grand average ERPs at 6 months in response to standard and deviant tones and white noise (Fig. 2a); the deviant-standard difference waveform (Fig. 2b); and the electrode locations used to calculate all auditory ERPs (Fig. 2c). ERPs to standard tones consist of the P150 followed by N250, and then the P300. ERPs to deviant tones and white noise represent a typical waveform consisting of a large and prolonged positive peak (merged P150 and early phase of P3a) (Kushnerenko et al., 2002). This resulted in the largest difference in amplitude of response to the frequently-presented standard tones and the less-frequent deviant tones occurring at around 300ms post stimulus onset. Topomaps show the development of the voltage distribution in seven 100ms bins from 100ms before stimulus onset to 500ms after stimulus onset showing an average of activity 50 ms around the peak (Fig 2d and 2e).

Fig. 3. shows the same information from the 12m visit: grand average ERPs following the standard, deviant and noise tones at 12m (Fig. 3a)); and the deviant-standard difference waveform (Fig. 3b)). ERPs to standard tones consist of the merged P150 and early phase of the P3 (or a P3 with a shorter latency), whereas ERPs to deviant tones represent a less merged double peak for the P150 and P3 in the same time window. This resulted in a deviant – standard difference wave peaking at a lower amplitude than at 6m at around 300ms post stimulus onset.

#### Visual task

Figure 4. Shows the grand average ERPs in response to fearful faces and neutral faces at 6m and 12m. The grand average waveforms clearly show P1, N290 and P4 components in response to faces at both ages. Topomaps show the development of the voltage distribution in seven 100ms bins from 100ms before stimulus onset to 500ms after stimulus onset showing an average of activity 50 ms +/- around the given value at both 6m and 12m.

### Analysis 1 – the development of associations between automatic neural sensitivity to auditory and visual stimuli

In Analysis 1 we examine how the relationship between neural sensitivity to auditory and visual stimuli develops between 6m and 12m of age.

#### 6-month data

First, we examined the associations between our auditory (the difference in amplitude of the P3 to standard and deviant tones) and visual (the difference in amplitude of the P1, N290 and P4 to neutral and fearful faces) neural sensitivity measures at 6m. Scatterplots illustrate the strength and direction of the correlation between each set of two variables (Fig. 6. a, c, f); The BFs above 1 indicate correlations for which the evidence from the current study is more likely under the hypothesis that there is a relationship between those variables in the population than not. That there is a positive relationship between the P3 auditory difference component and the P1 visual difference component in the population is nine times more likely from our evidence than not. That there is a negative relationship between the P3 auditory difference and the N290 visual difference component is 12 times more likely than not. That there is a positive relationship between the P3 auditory difference and the P4 visual difference component is five times more likely than not. Plots showing the prior and posterior distributions of the true population correlation show how evidence from the current study has updated the prior distribution (Fig. 6. b, d, f).

**Fig. 5.**
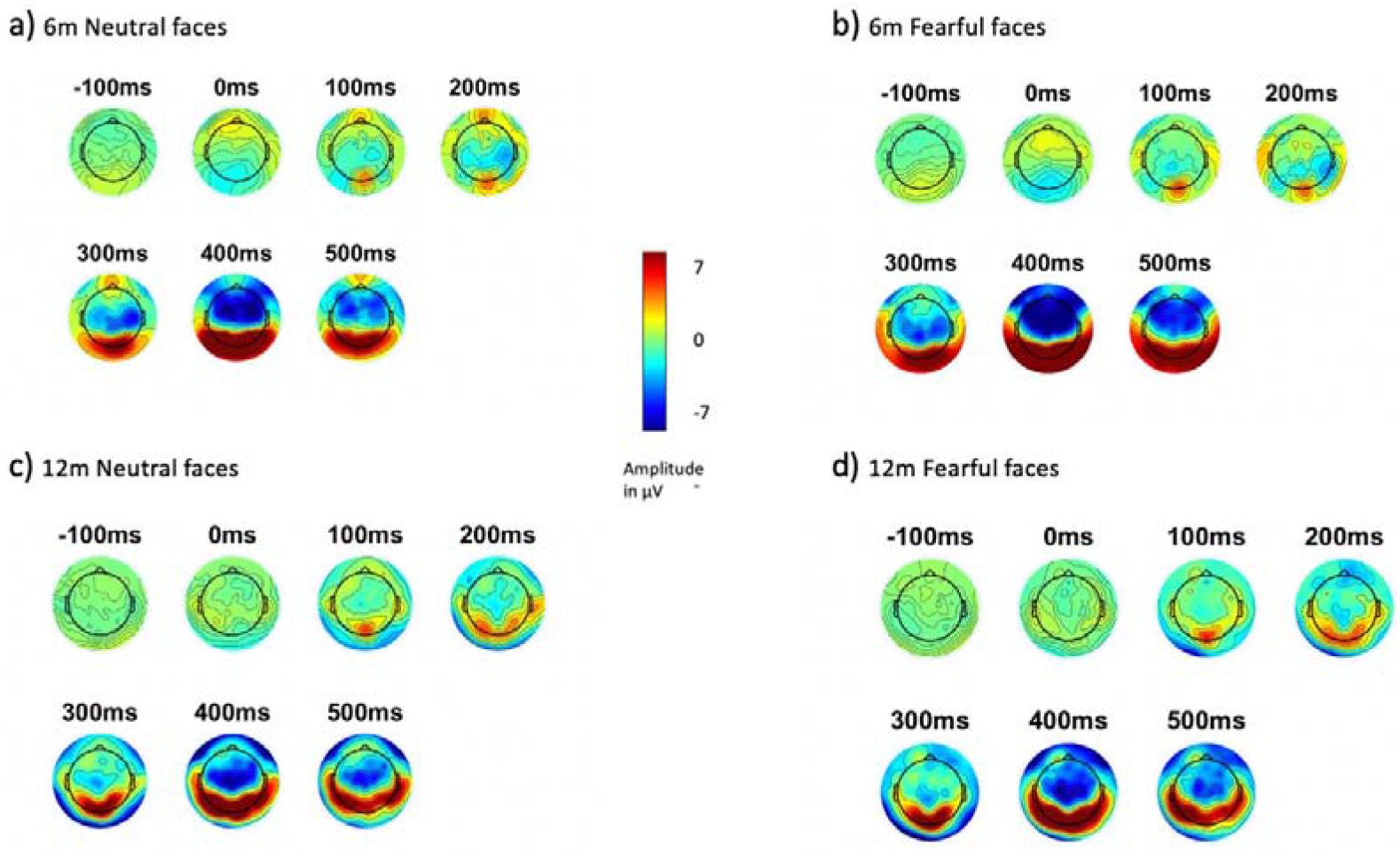
a and b) topomaps showing 6m response to neutral a) and fearful b) faces at 100ms intervals starting at 100ms before stimulus onset and ending 500ms post stimulus onset. c and d) topomaps showing 12m response to neutral c) and fearful d) faces at 100ms intervals starting at 100ms before stimulus onset and ending 500ms post stimulus onset. Each topoplot shows an average of activity 50 ms around the given value (i.e. −100ms shows the average from −150ms to −50ms). Topomaps were produced on data that was subject to channel interpolation outside of the main preprocessing

**Fig.6.**
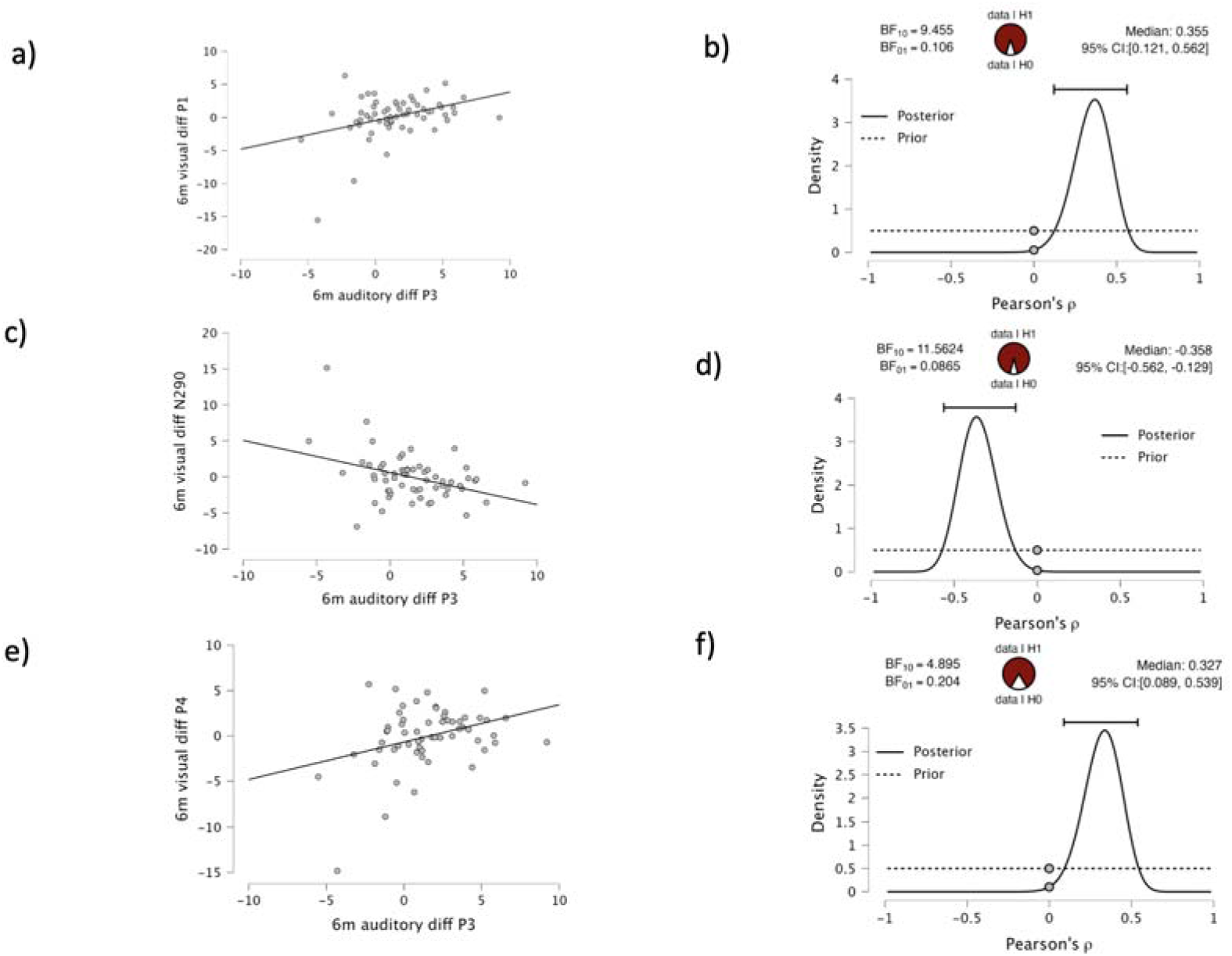
Correlations between measures of visual and auditory sensitivity at 6m: Scatterplots showing strength and direction of the Bayesian Pearson correlation between the two variables analysed (Fig.. 6…a, c, e); Plots showing the prior and posterior distributions of the true population correlation (Fig. 6. b, d, f). The BF is also presented graphically with the unit circle in the output. The shaded area corresponds to the evidence in favour of the alternative hypothesis (indicated in the graphic in Fig. 6 b) by “Data / H1”, and the unshaded area corresponds to evidence in favour of the null “Data \ H0”). The ratio of the shaded area to the unshaded area can be seen to be about 8:1 (6b), 12:1 (6d) 5:1 (6f), which is the value of BF10).

Table 1. shows the results of Bayesian Pearson correlations and Bayes Factor (BF) analyses. The Bayes Factor (BF10) for the relationship between the difference in the auditory P3 between standard and deviant tones and the visual P1 between neutral and fearful faces at 6m is 8. The BF for the negative relationship between the difference in the auditory P3 between standard and deviant tones and the visual N290 between neutral and fearful faces at 6m is 12. The BF for the relationship between the difference in the auditory P3 between standard and deviant tones and the visual P4 between neutral and fearful faces at 6m is 5. All results indicate there is moderate-strong evidence for rejecting the null hypothesis at 6m.

**Table 1.** Bayesian Pearson Correlations for neural auditory and visual sensitivity measures at 6m. Some of the Bayes Factors are exceptionally high: the BF for the association between visual diff P1 and visual diff P4 is 6.036 *10^6^.This example shows how a Bayesian analysis allows researchers to report a useful estimate of the exceptionally high strength of evidence (6 million to 1in favour of the alternative hypothesis) that would not be possible with a Pvalue

| Variable |  | 1. 6m auditory<br>diff_P3 | 2. 6m visual diff<br>P1 | 3. 6m visual diff<br>N290 | 4. 6m visual<br>diff P4 |
| --- | --- | --- | --- | --- | --- |
| **1. 6m auditory**<br>**diff P3** | n | — |  |  |  |
|  | Pearson's<br>r | — |  |  |  |
|  | BF□□ | — |  |  |  |
| **2.6m visual**<br>**diff P1** | n | 59 | — |  |  |
|  | Pearson's<br>r | 0.363 | — |  |  |
|  | BF□□ | 8.067 | — |  |  |
| **3. 6m visual**<br>**diff N290** | n | 61 | 60 | — |  |
|  | Pearson's<br>r | -0.373 * | -0.917 *** | — |  |
|  | BF□□ | 11.562 | 4.017e+21 | — |  |
| **4. 6m visual**<br>**diff P4** | n | 59 | 60 | 60 | — |
|  | Pearson's<br>r | 0.341 | 0.680 *** | -0.869 *** | — |
|  | BF□□ | 4.895 | 6.036e+6 | 2.254e+16 | — |

\* BF□□ > 10, \*\* BF□□ > 30, \*\*\* BF□□ > 100

#### 12-month data

Next, we conducted identical analyses on the 12-month data. Scatterplots illustrate the lack of a correlation between the two variables (Fig. 7. a, c, e); The BF01 above 1 indicates correlations for which the evidence from the current study is more likely under the null hypothesis that there is no relationship between those variables in the population. Plots showing the prior and posterior distributions of the true population correlation show how evidence from the current study has updated the prior distribution (Fig. 7. b, d, f).

**Fig. 7.**
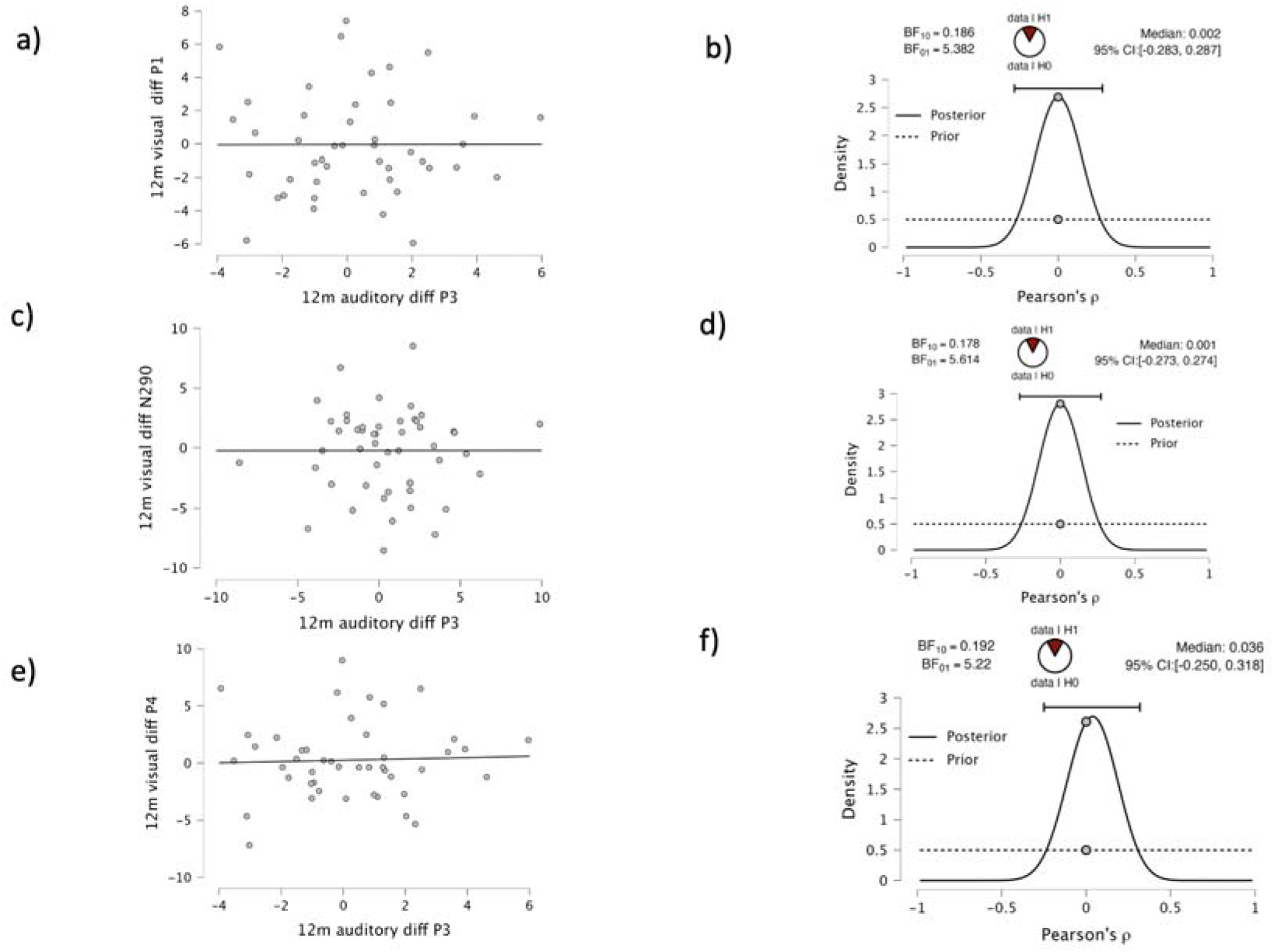
Correlations between measures of visual and auditory sensitivity at 12m: Bayesian correlation pairwise plots showing strength and direction of the Bayesian Pearson correlation between the two variables analysed (Figs. 7.a, c, e); Plots showing the prior and posterior distributions of the true population correlation (Figs. 7. b, d, f).

Table 2. shows the results of the Bayesian Pearson correlations and Bayes Factor (BF) analyses for the 12-month data. The BF10 (the likelihood of the data under the alternative compared to the null) for the relationship between the difference in the P3 component for standard and deviant tones and the difference in the P1 component for neutral and fearful faces at 12m is 0.2. The BF10 for the relationship between the difference in the P3 component for standard and deviant tones and the difference in the N290 component for neutral and fearful faces is 0.2. The BF10 for the relationship between the difference in the P3 component for standard and deviant tones and the difference in the P4 component for neutral and fearful faces is 0.2. This can be interpreted as our evidence being “moderately” more likely under the null hypothesis that these measures of neural sensitivity are not correlated in the population at 12m.

**Table 2.** Bayesian Pearson Correlations for neural auditory and visual sensitivity measures at 12m

| Variable | 1. 12m<br>auditory<br>diff P3 | 2. 12m<br>visual<br>diff P1 | 3. 12m<br>visual<br>diff N290 | 4. 12m<br>visual<br>diff P4 |
| --- | --- | --- | --- | --- |
| **1. 12m**<br>**auditory**<br>**diff P3** |  |  |  |  |
| n | — |  |  |  |
| Pearson's<br>r | — |  |  |  |
| BF<sub>10</sub> | — |  |  |  |
| **2. 12m**<br>**visual diff**<br>**P1** |  |  |  |  |
| n | 460 | — |  |  |
| Pearson's<br>r | -0.021 | — |  |  |
| BF<sub>10</sub> | 0.186 | — |  |  |
| **3. 12m**<br>**visual diff**<br>**N290** |  |  |  |  |
| n | 49 | 47 | — |  |
| Pearson's<br>r | 0.001 | -0.943 *** | — |  |
| BF<sub>10</sub> | 0.178 | 6.738e+19 | — |  |
| **4. 12m**<br>**visual diff**<br>**P4** |  |  |  |  |
| n | 46 | 47 | 47 | — |
| Pearson's<br>r | 0.035 | 0.766 *** | -0.865 *** | — |
| BF<sub>10</sub> | 0.189 | 3.431e+7 | 1.498e+12 | — |

\* BF<sub>10</sub> > 10, \*\* BF<sub>10</sub> > 30, \*\*\* BF<sub>10</sub> > 100

Overall, the results from Analysis 1 indicate that at 6-months of age, indices of neural sensitivity to auditory information (P3) and visual information (P1 and P4) are correlated positively, whereas the auditory difference P3 is negatively correlated with the visual difference N290. However, by 12-months of age any association between these measures has disappeared.

### Analysis 2 – the association between autonomic arousal and visual and auditory neural sensitivity

For the second analysis, we examine how autonomic arousal related to neural sensitivity at 6m and 12m. We operationalised autonomic arousal as heart rate (HR) in beats per minute (BPM) averaged across the recording. Neural sensitivity on the auditory task was operationalised as the amplitude difference in the P3 components in response to standard and deviant tones. Neural sensitivity on the visual task was operationalised as the amplitude difference in the P1, N290 and P4 components in response to fearful and neutral faces. At 6m, HR correlated negatively with the auditory P3 difference and the visual P4 difference. However, HR correlated positively with the visual N290 difference. No correlation at a statistically significant level was found between HR and the difference between facial expressions in the P1 component. Follow-up analyses showed that autonomic arousal associated with a larger negative-going N290 in response to fearful but not neutral faces.

Scatterplots illustrate the strength and direction of the correlation between each set of two variables (Fig. 8. a, c, e). Our evidence shows that a negative relationship between average HR in BPM over the entire recording and the difference in the P3 component for standard and deviant tones and the P4 component for neutral and fearful faces is twice and four times respectively as likely as no relationship in the population. The BF10 for the positive relationship between average HR in BPM over the entire recording and the difference in the N290 component to neutral and fearful faces shows that a positive correlation is 12 times more likely in the population than not.

**Fig. 8.**
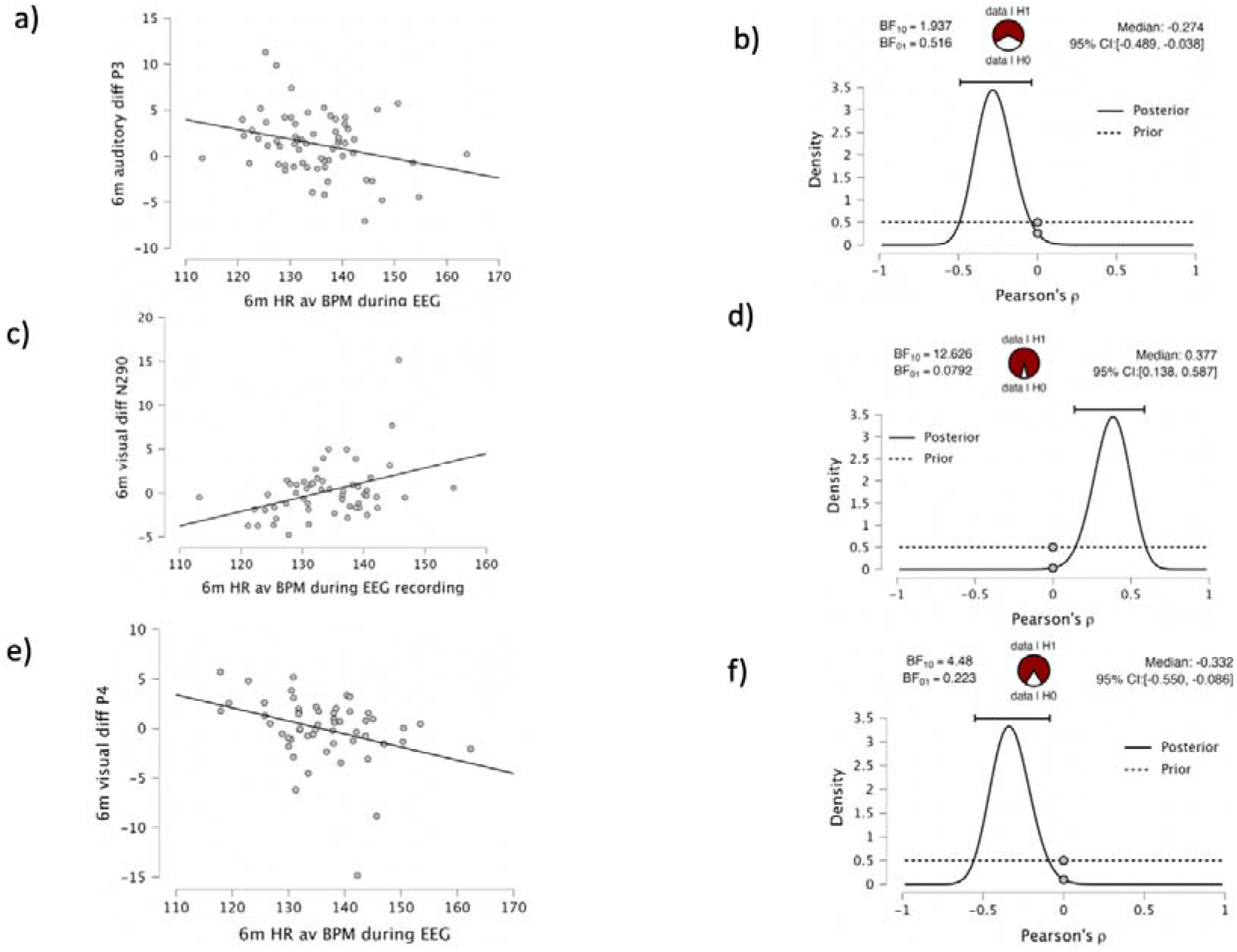
Correlations between measures of HR and visual and auditory sensitivity at 6m: Bayesian correlation pairwise plots showing strength and direction of the Bayesian Pearson correlation between the two variables analysed (Figs. 8. a, c, e); Plots showing the prior and posterior distributions of the true population correlation (Figs. 8. b, d, f).

Plots showing the prior and posterior distributions of the true population correlation show how evidence from the current study has updated the prior distribution (fig. 8. b, d, f)

Table 3 shows the Bayesian Pearson Correlations for the autonomic and neural measures.

**Table 3.** Bayesian Pearson Correlations for autonomic and neural measures at 6m

| Bayesian Pearson Correlations |  | 1. BPM 6m | 2. 6m auditory diff P3 | 3. 6m visual diff N290 | 4. 6m visual diff P4 |
| --- | --- | --- | --- | --- | --- |
| 1. BPM 6m | n | — |  |  |  |
|  | Pearson's r | — |  |  |  |
|  | BF□□ | — |  |  |  |
| 2. 6m auditory diff P3 | n | 62 | — |  |  |
|  | Pearson's | -0.287 | — |  |  |

| Variable | 1. BPM 6m | 2. 6m auditory diff P3 | 3. 6m visual diff N290 | 4. 6m visual diff P4 |
| --- | --- | --- | --- | --- |
| r |  |  |  |  |
| BF$\square\square$ | 1.933 | — | | |
| **3. 6m visual diff N290** |  |  |  |  |
| n | 55 | 61 | — |  |
| Pearson's r | 0.394* | -0.371* | — |  |
| BF$\square\square$ | 12.626 | 10.900 | — | |
| **4. 6m visual diff P4** |  |  |  |  |
| n | 53 | 59 | 60 | — |
| Pearson's r | -0.341 | 0.254 | -0.869*** | — |
| BF$\square\square$ | 3.564 | 1.025 | 2.254e+16 | — |

\* BF $\square\square$ > 10, \*\* BF $\square\square$ > 30, \*\*\* BF $\square\square$ > 100

Scatterplots in Figure 9. show the strength and direction of the correlation between autonomic arousal and the N290 to fearful faces at 6m (Fig. 9 a) and 12m (Fig. 9 b). Plots showing how the evidence has updated the prior distributions for the relationship at 6m to be 3 times more likely than not (Fig. 9. C) and at 12 months, that there is no relationship between the variables being three times more likely than that there is.

**Fig. 9.**
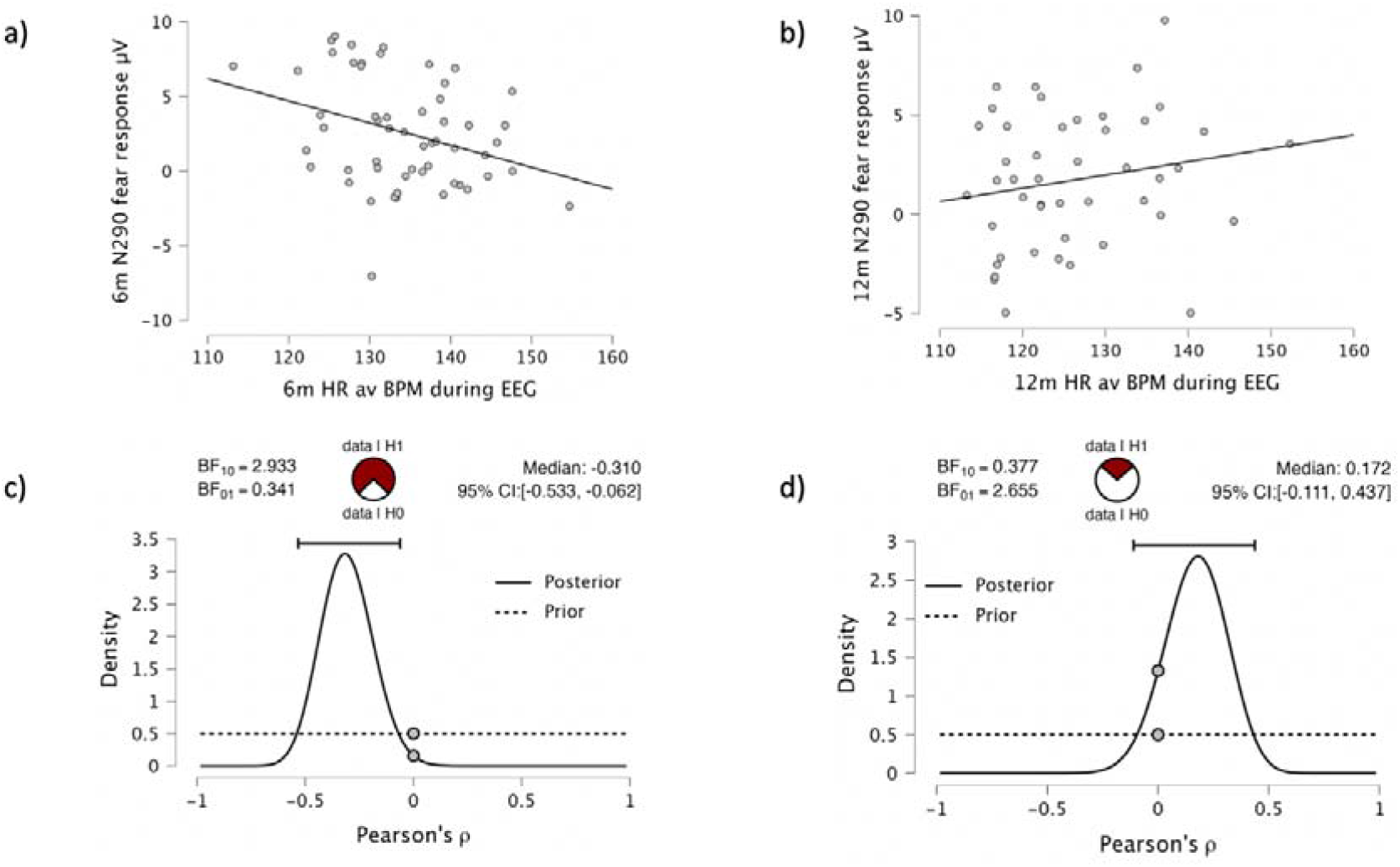
Correlations between measures of HR and visual N290 response to fearful faces at 6m and 12m: Bayesian correlation pairwise plots showing strength and direction of the Bayesian Pearson correlation between the amplitude of the 6m N290 to fear and HR at 6m (Fig. 9. a) and the amplitude of the 12m N290 to fear and HR at 12m (Fig. 9. b); Plots showing the prior and posterior distributions of the true population correlation (Figs.9. c, d).

**Fig. 10.**
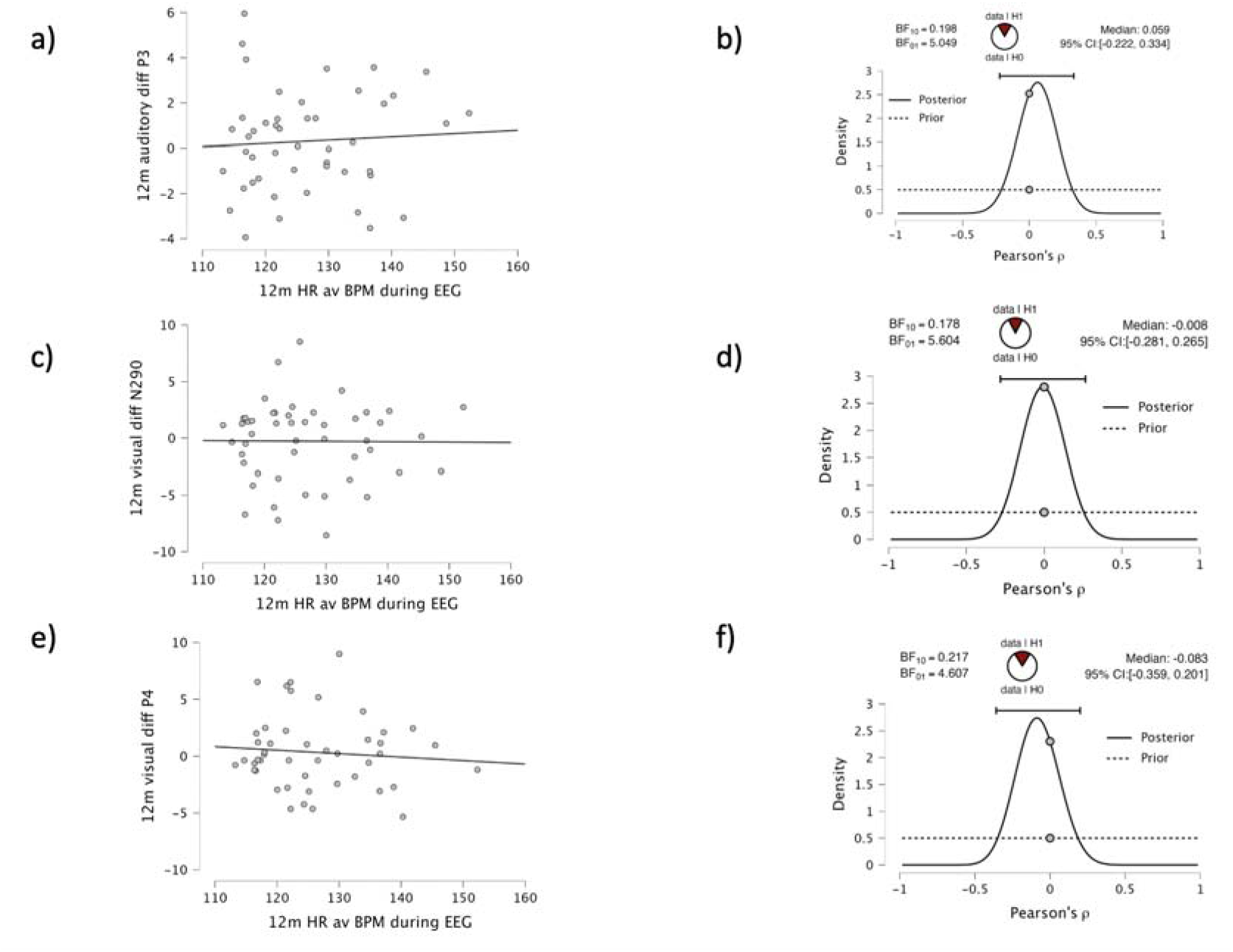
Correlations between measures of HR and visual and auditory sensitivity at 12m: Bayesian correlation pairwise plots showing strength and direction of the Bayesian Pearson correlation between the two variables analysed (Figs. 10.a, c, e); Plots showing the prior and posterior distributions of the true population correlation (Figs. 10. b, d,f))

Table 4. shows the Bayesian Pearson Correlations for autonomic arousal and N290 in response to fear at 6m and 12m.

**Table 4.**
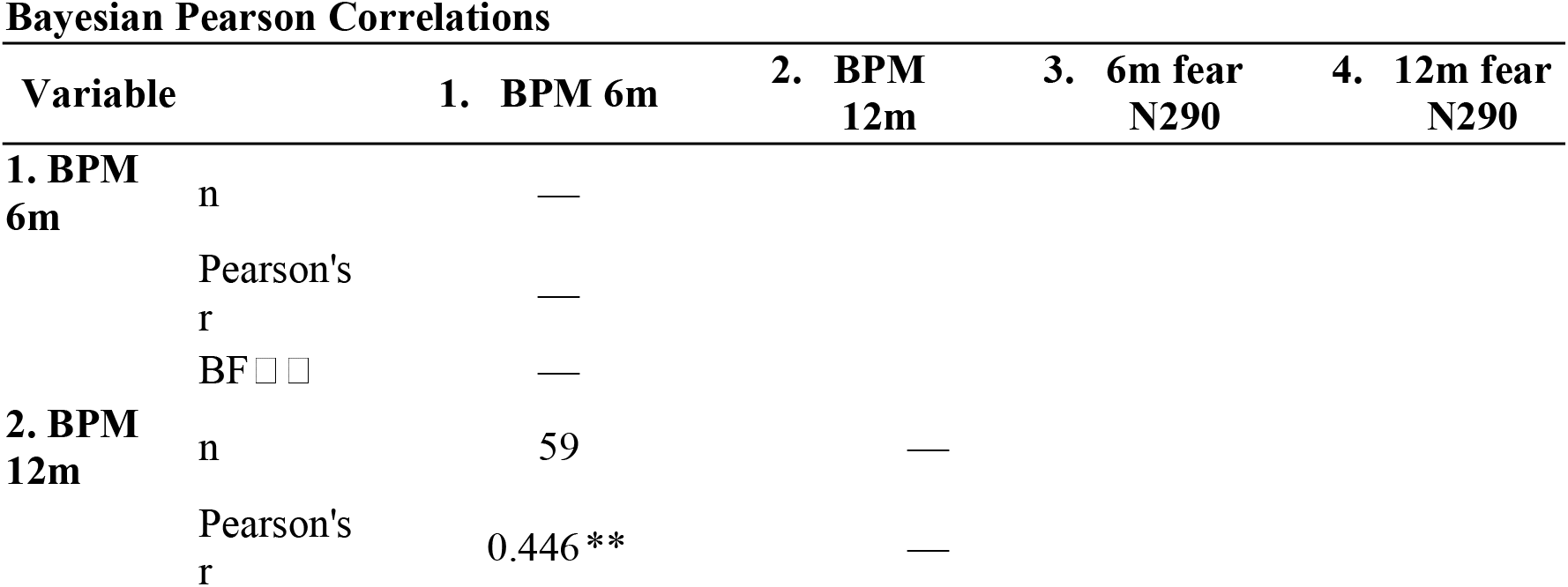
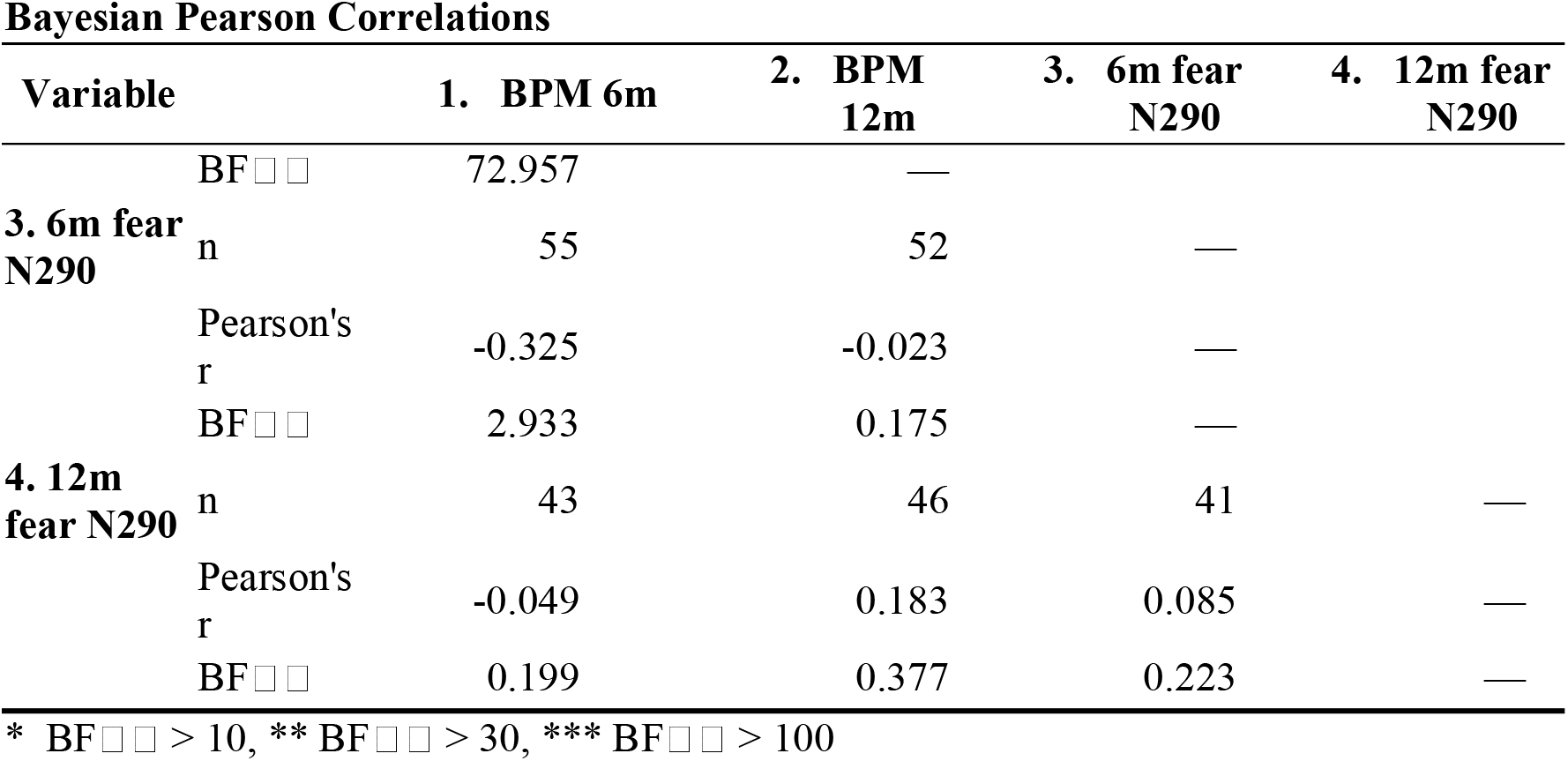
Bayesian Pearson Correlations for measures of autonomic arousal and N290 response to fearful faces at 6m and 12m

**Table 5.** Bayesian Pearson Correlations for autonomic and neural measures at 12m

| Variable |  | 1. BPM<br>12m | 2. 12m<br>auditory<br>diff P3 | 3. 12m visual<br>diff P1 | 4. 12m visual<br>diff N290 | 5. 12m<br>visual<br>diff P4 |
| --- | --- | --- | --- | --- | --- | --- |
| **1. BPM**<br>**12m** | n | — |  |  |  |  |
|  | Pearson's<br>r | — |  |  |  |  |
| | BF$\square\square$ | — | | | | |
| **2. 12m**<br>**auditory**<br>**diff P3** | n | 54 | — |  |  |  |
|  | Pearson's<br>r | 0.045 | — |  |  |  |
| | BF$\square\square$ | 0.178 | — | | | |
| **3. 12m**<br>**visual diff**<br>**P1** | n | 46 | 46 | — |  |  |
|  | Pearson's<br>r | -0.086 | -0.021 | — |  |  |
| | BF$\square\square$ | 0.215 | 0.186 | — | | |
| **4. 12m**<br>**visual diff**<br>**N290** | n | 49 | 49 | 47 | — |  |
|  | Pearson's<br>r | -0.009 | 0.001 | -0.943 *** | — |  |
| | BF$\square\square$ | 0.178 | 0.178 | 6.738e+19 | — | |
| **5. 12m**<br>**visual diff**<br>**P4** | n | 46 | 46 | 47 | 47 | — |
|  | Pearson's<br>r | -0.089 | 0.035 | 0.766 *** | -0.865 *** | — |
| | BF$\square\square$ | 0.217 | 0.189 | 3.431e+7 | 1.498e+12 | — |
| * BF$\square\square$ > 10, ** BF$\square\square$ > 30, *** BF$\square\square$ > 100 | | | | | | |

These results indicate that at 6m there is a negative relationship between physiological arousal and neural sensitivity as operationalised in this study at the auditory P3 component and visual P4 component. However, individuals with higher autonomic arousal during the EEG recording session, responded with a larger difference in amplitude between fearful and neutral faces at the N290 component.

Next, we repeated an identical analysis based on the 12m data. Scatterplots illustrate the strength and direction of the correlation between each of the sets of two variables (Fig. 9. a, c, e); Our evidence shows that no relationship between average HR in BPM over the entire recording and the difference in the P3 component for standard and deviant tones and the, N290 and P4 components for neutral and fearful faces is five, six and four times more likely, respectively, than there being a relationship in the population. Plots showing the prior and posterior distributions of the true population correlation show how evidence from the current study has updated the prior distribution (fig. 9. b, d,f).

Overall, the results from Analysis 2 indicate that at 6m, higher physiological arousal associated with decreased neural sensitivity in the auditory domain (specifically, a larger difference between the amplitude of responses to standard of deviant tones at the P3 component) and the visual domain (a larger difference between the amplitude of responses to fearful and neutral faces at the P4 component). However, higher physiological arousal also associated with increased neural sensitivity as measured by the difference between the amplitude of responses to fearful and neutral faces at the N290 component. Autonomic arousal did not correlate with any measures of neural sensitivity at 12m.

## Discussion

We used ERP paradigms to measure auditory and visual sensitivity in infants at 6 and 12m while concurrently measuring inter-individual differences in autonomic arousal. Our results had two main features of interest. The first was that at 6 months, neural sensitivity - indexed by detection of difference - correlated across auditory and visual modalities. Specifically, we observed positive associations between the difference in response amplitudes of the P1 and P4 between the neutral and fearful conditions in the visual paradigm and the difference in response amplitudes of the P3 component between the standard and deviant conditions in the auditory paradigm. We also found a negative association between the difference in response amplitude of the N290 between neutral and fearful conditions and the difference in response amplitudes of the P3 between the standard and deviant conditions in the auditory paradigm. The same associations were not present at 12-months. Second, at 6m, infants’ autonomic arousal negatively correlated with the auditory difference P3 and the visual difference P4 but positively correlated with the visual difference N290. Any association between autonomic arousal and neural sensitivity disappeared at 12m despite the 6m and 12m EEG measures having comparable levels of noise and variability. We shall discuss these two main findings in turn.

Topomaps of our results show that different cortical regions are being activated in response to the visual and auditory stimuli. While this implies specialisation of cortical areas for visual (occipital) and auditory (temporal) perception, our measures of neural sensitivity (indexed as difference-detection between conditions) nevertheless correlate at 6m. This implies a domain general level of sensitivity, in terms of the early stages of visual and auditory processing which has been considered evolutionarily adaptive – to facilitate making novel and serendipitous associations with environmental cues in an uncertain environment (Chiappe & MacDonald, 2005). This early universal sensitivity is corroborated by examinations of the relationship between neural responses to visual and auditory stimuli in the same infants during studies into the development of audiovisual speech integration: Despite the different modalities being presented simultaneously, most evidence suggests that young infants process signals from the two modalities separately and do not integrate them into a single percept until at least 4.5-months old (Bristow et al., 2009; Desjardins & Werker, 2004; Nardini et al., 2010). This evidence corroborates theories on perceptual narrowing in language development (Scott et al., 2016). Our auditory stimuli consisted of non-speech sounds, so would not normally co-occur with faces. However, the initial covariance of sensitivity to auditory and visual stimuli is in line with the intersensory integration view (Birch & Lefford, 1963, 1967), which posits that the different sensory systems do not communicate with one another at birth and, at least in humans, intersensory liaisons are not formed until the first months or even years of life. Near-infrared spectroscopy (NIRS) showed that unlike adults, 5m-old infants recruit adjacent but non-overlapping regions in the left dorsal prefrontal cortex when they process eye contact and own name and do not seem to recruit a common prefrontal region when processing communicative signals of different modalities (Grossmann et al., 2010).

Another explanation for the early covariance of difference-detection in visual and auditory domains may lie in the ontogeny of face-processing. Some argue that there is an innate cortical module specifically dedicated to face processing (Farah et al., 2000). However, in very young infants, Johnson argued it was sub-cortical orienting involving the amygdala that modulates activity in face-sensitive cortical regions before the arrival of visual information through the cortical route (Johnson, 2005). The pathway is maximally sensitive to low-spatial-frequency (LSF) aspects of faces, which selectively differentiates expressions such as fear with wider eyes and open mouths. This sensitivity to LSF aspects of faces could explain the early association between some components indexing visual and auditory neural sensitivity in this study. This can be seen in the functional specificity of the different ERP components and the direction in which they correlate. In adults, the longer-latency visual ERP components, greater than 400ms after stimulus onset, are thought to indicate recognition of facial identity (Barrett et al., 1988; Barrett & Rugg, 1989; Eimer, 2000; Itier & Taylor, 2004) and/or retrieval of semantic information related to faces (Paller et al., 2000). However, the earlier N170 is thought to be related to stages of structural encoding of the physical information in faces, with some studies suggesting that it may only reflect eye detection (Bentin et al., 1996) as opposed to encoding of the entire configuration of facial features (Eimer, 2000). In addition, the N170 component can be unaffected by any emotional expression, supporting the hypothesis that structural encoding and expression analysis are independent processes (Eimer et al., 2003). In infants, there is evidence that the adult N170 is preceded by the N290 and P4 components. However, before 12 months of age, the P4 (unlike the N290) component does not seem to be face-specific (Halit et al., 2003b). Furthermore, the amplitude of the P4 was not sensitive to the difference between face and visual-noise stimuli in 3-month-olds, while the amplitude of the N290 displayed a huge difference (Halit et al., 2004). In addition, the P1 is an obligatory visual component indexing low-level sensory processing and is not face-specific but associated with differences in low-level visual features that exist between face and non-face stimuli (Conte et al., 2020). Moreover, Cohen and Cashon (2001) report a developmental shift at 7m from featural to configural processing of faces. They suggest that before the age of 7 months, infants process specific features of complex objects but after the age of 7 months they are able to integrate those features into a whole object (Cohen & Cashon, 2001; Conte et al., 2020). Taking the above evidence into account, it is proposed that the difference between the amplitude of the early P1 and P4 components to the fearful and neutral faces at 6m may be explained by the encoding of the lower-level, perceptual information in the isolated features of the faces such as larger eyes and open or down-turned mouths in the fearful category (Halit et al., 2003b). Adult studies have seen larger amplitude N170 components evoked to open as opposed to closed mouths (Puce et al., 2007; Wheaton et al., 2001). The P1 and P4, components are more likely to be affected by the spatial differences of fearful as opposed to neutral faces detected by exogenous attention, whereas the difference in amplitude of the N290 evoked by the two visual conditions may be partially explained by the recruitment of greater top-down, pre-frontally mediated processing, which would not associate positively with indexes of exogenous perception – the auditory P3 and the visual P1 and P4. This differentiation between exogenous perception of low level sensory features and the more experience-mediated endogenous processing corresponds with the proposition that two different mechanisms underlie the auditory tracking of the speech envelope: one derived from the intrinsic oscillatory properties of auditory regions; the other induced by top-down signals coming from other non-auditory regions of the brain (Rimmele et al., 2018). Under nonspeech listening conditions, the intrinsic auditory mechanism dominates (Assaneo et al., 2019), which corresponds with the automatic change detection in the processing of non-sematic, lower-level features of non-speech sounds in this study. In contrast, at 12-months, a developmental shift may have occurred whereby infants have developed a more adult-like processing mechanism which integrates features to process a face as special category of object. Perhaps at 12m in this sample we have captured a transition from bottom-up, stimulus-driven processing to more top-down processing - based on experience - which no longer correlates with the auditory mismatch response which is thought to be automatic and independent of voluntary attention (Cheour et al., 2010; Háden et al., 2016; Wanrooij et al., 2014).

The dissociation of visual and auditory sensitivity by 12m could also be due to the differential development in the two modalities. There is ample evidence that very early in development audio and visual development rates differ. Differential onset of the functioning of sensory systems results in relative independence among emerging systems, thereby reducing competition which helps regulate subsequent neurogenesis and functioning (Turkewitz & Kenny, 1982). Synaptogenesis and synapse elimination occurs at different rates in different cortical regions in humans. Synaptic density in the auditory cortex is maximal at 3-months of age and synaptic elimination ends at around 12-years, whereas synaptic density in the visual cortex is maximal between 9 and 15-months and synaptic elimination ends at around late adolescence (Huttenlocher and Dabholkar, 1997). Myelination begins earlier in the occipital lobe than in the temporal lobe after birth (Yakolev and Lecours, 1967). Complexity measures, such as multiscale entropy (MSE) (Costa et al., 2002) can index maturational changes in brain function. While EEG signal complexity increased from one month to 5 years of age in response to auditory and visual stimulation, infants’ signal complexity for the visual condition was greater than auditory signal complexity, whereas adults showed the same level of complexity to both types of stimuli. The differential rates of complexity change may reflect a combination of innate and experiential factors on the structure and function of the two sensory systems (Lippé et al., 2009).

The second branch of our findings on the domain generality of ES was that higher heart rate (HR) (measured in BPM averaged across the entire EEG recording) was associated with a larger difference in response amplitude of the visual N290 component to fearful and neutral faces and a smaller difference in the response amplitudes of the visual P4 component and the auditory P3 component in the 6-month-old infants. While HR correlated between 6m and 12m, any associations between autonomic arousal and automatic neural sensitivity had disappeared by 12-months despite equal amounts of noise and variability at the two time-points (see error bars in Figures 2. 3. and 4.) In the same auditory change-detection paradigm as used in this study, while responses to large acoustic contrasts (bursts of white noise) evoked large P3 responses (indexing exogenous, stimulus-driven orienting or distractibility) in all 5-7 year-old children regardless of HR, children with high autonomic arousal also showed a larger P3 component in response to small acoustic contrasts (500Hz-750Hz) (Wass et al., 2019). It was hypothesized that in trials with high HR, the overall brain excitability was higher and therefore more prone to involuntary attention. Thus, even small acoustic contrasts (frequency deviant) could potentially elicit a P3-like response. Therefore, for this study we hypothesized that higher HR would associate with greater neural sensitivity indexed by a larger difference in the amplitude of response to the two conditions in the visual and auditory paradigms. However, we only found a larger difference in the amplitude of response to fearful and neutral faces at one component – the N290. This finding may be explained by the follow-up analyses, which showed that high HR correlated with larger N290 (but not P1 or P4) responses to fearful faces. The differential response at the different components will be addressed below, but the larger N290 to fearful faces could index greater exogenous attention to salient environmental stimuli. Heightened autonomic arousal is an index of sympathetic nervous system (SNS) activity which is involved in quick response mobilising (‘fight or flight’) (Cacioppo et al., 2000) and as such is associated with hypervigilance and sensory reactivity to environmental stimuli (Cheung & Porges, 2013). Relatedly, prior research has found associations between sensory-reactivity and emotional-face processing in children. Projections from the amygdala (part of the neural system responsive to threat (Tovote et al., 2015)) to the occipital cortex may serve to enhance the processing of visually salient stimuli, including facial expressions of emotion (Eimer et al., 2003) and especially fearful expressions (Morris et al., 2002).

The difference in the direction of the correlations between components indexing neural sensitivity and measures of physiological arousal may again be explained by the functional specificity of the components. Whereas, as mentioned above, there is evidence of a difference in response amplitude of the N290 between face and non-face visual stimuli suggesting the N290 is face-specific (Halit et al., 2003b), the P1 is an obligatory visual component indexing low-level sensory processing and is not face-specific (Conte et al., 2020) and the P4 is thought to reflect structural processing of faces in infants, (Porter et al., 2021) but also does not seem to be face-specific (Halit et al., 2003b). The fact that detection of difference at these visual components correlates positively with detection of difference in the P3 auditory component may be because all three index stimulus-driven low-level perception of the sensory properties of the stimuli in the two modalities. For the same reason, the direction of the correlation between HR and the amplitude of the difference between these components is the same - slower heart rate was associated with greater neural sensitivity in terms of perception of difference between conditions in stimuli for the auditory P3 and the visual P4 component.

This study set out to answer the question of whether environmental sensitivity is domain-general or domain-specific. Evidence presented here suggests that neural sensitivity, in terms of automatic exogenous perception of salient stimuli, is related in different modalities at 6m and associates with autonomic arousal, but then follows different developmental trajectories. An initial domain generality of sensory sensitivity may confer advantages in that an organism is initially equipped to respond to any environmental stimuli, and then gradually develops an expertise for the stimuli to which it is predominantly exposed.

Further research aimed at understanding the relationship between different sensory modalities and how these associate with autonomic states would improve our understanding of the autonomic and neural mechanisms associated with a hypothesised common factor of sensitivity. This would contribute towards finding an easily measurable phenotype for environmental sensitivity. Being able to measure individual differences in environmental sensitivity would facilitate the provision of appropriate levels of stimulation in developmental environments.

This study was funded by the Economic and Social Research Council (ESRC)

## Supplementary materials

In the automatic artefact correction algorithm, artefacts are detected from dRR series, which is a time series consisting of differences between successive RR intervals. The dRR series provides a robust way to separate ectopic and misplaced beats from the normal sinus rhythm. To separate ectopic and normal beats, time varying threshold (Th) is used. To ensure adaptation to different HRV levels, the threshold is estimated from the time varying distribution of the dRR series. For each beat, quartile deviation of the 90 surrounding beats is calculated and multiplied by factor 5.2. Beats within this range cover 99.95% of all beats if RR series is normally distributed. However, RR interval series is not often normally distributed, and thus, also some of the normal beats exceed the threshold. Therefore, decision algorithm is needed to detect artefact beats. Ectopic beats form negative-positive-negative (NPN) or positive-negative-positive (PNP) patterns to the dRR series. Similarly long beats form positive-negative (PN) and short beats negative-positive (NP) patterns to the dRR series. Only dRR segments containing these patterns are classified as artefact beats. Missed or extra beats are detected by comparing current RR value with median of the surrounding 10 RR interval values (medRR). A missed beat is detected if current RR interval (RR(i)) satisfies condition RR(i) 2 - medRR(i) < 2T h (1) and an extra beat is detected if two successive RR intervals (RR(i) and RR(i+1)) satisfy condition |RR(i) + RR(i + 1) - medRR(i)| < 2T h. (2) Detected ectopic beats are corrected by replacing corrupted RR times by interpolated RR values. Similarly too long and short beats are corrected by interpolating new values to the RR time series. Missed beats are corrected by adding new R-wave occurrence time and extra beats are simply corrected by removing extra R-wave detection and recalculating RR interval series

